# Bacterial exposure shapes the trajectory of immune ageing

**DOI:** 10.1101/2025.10.10.681722

**Authors:** Erik Feldman, Michal Hrúzik, Holden T. Maecker, Alain Roulet, Benjamin Lelouvier, Neta Milman, Martin Lukačišin, Shai S. Shen-Orr

**Affiliations:** Department of Immunology, Faculty of Medicine, Technion - Israel Institute of Technology, 1 Efron St, 35254, Haifa, Israel; Department of Genetics, Faculty of Natural Sciences, Comenius University, Ilkovičova 6, 84104, Bratislava, Slovakia; The Human Immune Monitoring Center, Stanford University School of Medicine, 299 Campus Drive, Stanford, California, 94305, USA; Vaiomer, 516 rue Pierre et Marie Curie, 31670, Labège, France

## Abstract

The immune system is uniquely capable of both regenerating and ageing the rest of the organism based on its own state^1^. Understanding the driving forces behind alterations in the immune system with age^2^ has thus potential to lead to interventions against ageing in general^3^. However, the factors driving immune ageing remain largely unknown^4^. Here we show that alterations in response to bacterial exposure are a key driving force of immune ageing. Studying a longitudinal cohort with their immune system characterised in high-dimension^2^, we devised a phenotype-driven gene co-regulation analysis through which we unravel the immune-ageing role for genes regulated by miRNAs previously reported to cause gut leakiness^5^. Hypothesizing increased bacterial exposure as a driver of immune ageing, we estimated the exposure of each individual’s immune system to bacteria through analysing the residual bacterial DNA content in their immune cells^6^. We found that the total bacterial load is unrelated to the subject’s chronological or immune age, but predicts the magnitude of change in immune age over the next year. Finally, we rationalise previously observed alterations in cellular composition of the immune system associated with ageing^2^ by finding that the immune cell subtypes most increasing in their frequency are highly enriched in expressing bacterial response genes. Our results thus earmark bacterial exposure as the candidate driver to be counteracted in anti-ageing interventions and the immune cell residual bacterial DNA content as a biomarker for the immune systems most at risk of ageing. Overall, these results offer a new paradigm for investigating gut health in the context of immune ageing^7,8^.

## Main

Ageing of biological systems is commonly understood as a time-dependent decline in their essential physiological functions^9^. Ageing in humans has been recently characterised in a variety of molecular and organismal readouts^10–13^. Connection of molecular readouts with phenotypes such as morbidity or mortality allowed for creation of a number of phenotypic ageing metrics^2,14–19^ ‒ their joint characteristic is that they replace the chronological age of a subject with a biological age that marks the progression of an individual on a directional trajectory leading to the phenotype in question, such as death. Through such phenotypic ageing metrics, it has been shown that different organs in the same organism can age differentially and thus a given individual can possess multiple biological ‘ages’ pertaining to different components of the body^20–23^.

The immune system, however, plays a special role in the aging of the entire organism^3^. Being involved in maintaining organismal homeostasis, it is bi-directionaly involved with all tissues and organs, as evidenced by parabiosis experiments showing that an aged immune system transplanted into a young animal ages its peripheral organs, just as the immune system from the young animal rejuvenates an aged individual^1^. The immune system thus warrants special attention in investigating biological ageing in general. We and others have recently devised immune ageing metrics and quantitatively characterised phenotypic changes associated with the ageing of the immune system^2,24–26^. While individuals age at different rates, immune ageing follows stereotyped patterns, such as coordinated changes in the frequency of specific immune cell subtypes^2^. These phenotypic descriptions are a necessary pre-condition for investigating the drivers of immune ageing; however, the progress in identifying drivers has been piecemeal^27^.

In this work we develop a statistical method to characterise gene regulatory changes during immune ageing and use this framework to generate the hypothesis that it is an increased exposure of the immune system to bacteria, potentially through increased gut permeability, that drives immune ageing. We then experimentally corroborate this hypothesis by showing that bacterial DNA present in the immune cells quantitatively predicts ageing of the immune system in one year following the original sampling. Finally, we conceptually link the previous phenotypic description of changes in immune cell frequencies during immune ageing to this driver through finding that cell subtypes previously found to increase during immune ageing are those expressing genes involved in responding to bacterial presence.

### Gene-regulatory remodelling during immune ageing

In the search for drivers of systemic changes in the ageing immune system, we reasoned that it would be advantageous to quantify not only changes in gene expression levels during immune ageing^2^, but also changes in gene regulatory networks. To disentangle such immune age-associated remodelling of gene regulation, we devised an approach that we term phenotype-driven gene correlation network analysis (pCNA). Briefly, the gene expression measurements for individual samples are first ordered according to a quantitative phenotype, such as immune age, followed by quantification of pairwise gene-gene correlations within a sliding window (**Fig. 1a**). The underlying principle is that since the regulatory relationships themselves might change and in that way bring about the change in phenotype, we divide the entire set into multiple subsets stratified by the phenotype in question (**Fig. 1a**). This stands in contrast to the commonly performed gene correlation network analysis^28^, where the gene-gene wiring is assumed to be constant and the phenotypes are associated with changes in the expression level of the clusters of co-regulated genes, rather than with changes in their wiring.

**Figure 1:**
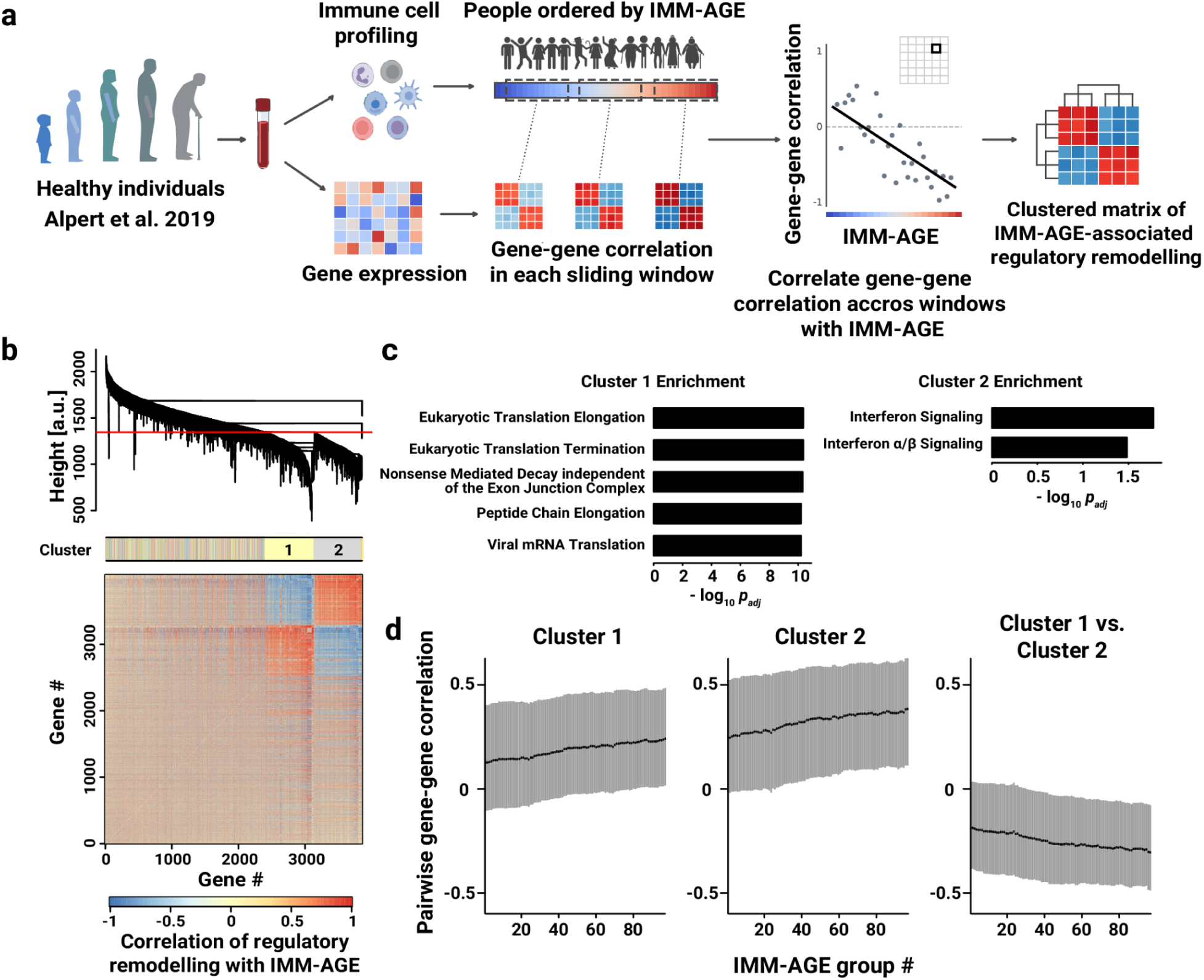
Phenotype-driven gene correlation network analysis (pCNA) of immune ageing. **a**, Schematic of pCNA on microarray samples from Stanford Elison cohort. Samples are ordered according to the donor’s immune age as determined from immune cell proportions and pairwise gene expression correlations calculated within a sliding window. Gene-gene correlations are then correlated with IMM-AGE across all window positions. **b**, Heatmap of correlations between IMM-AGE and gene-gene correlations after hierarchical clustering. Colour bar shows cluster identity for genes according to the thresholding of the distance tree shown at the top. **c**, Functional enrichment of genes in two largest clusters from (**b.**) using Reactome gene ontology. Top five enriched terms with *p_adj_* < 0.05 are shown for each cluster. **D**, Boxplots of pairwise gene-gene correlations for all sliding-window positions for genes in the two largest gene clusters. The sets of boxplots show gene-gene correlations within the gene sets (left and middle) or between the gene sets (right); black line ‒ median, grey area ‒ interquartile range.

We performed pCNA on the gene expression microarray data from the SELA (Stanford Elison Longitudinal Aging) cohort, a longitudinal cohort of healthy people previously characterised using multiple high-throughput modalities^2,29,30^, and currently running for over 15 years. Specifically, we ordered the samples by their inferred IMM-AGE (ref. 4; *Methods*), a metric of immune age determined by diffusion analysis of relative frequencies of immune cell subsets, shown to be predictive of all-cause mortality and other clinical phenotypes even after regressing out the most common predictors^2^. Performing pCNA, we discovered two gene modules that with increasing immune age are more tightly co-regulated within each cluster, but increasingly anti-coregulated between the two clusters (**Fig. 1b**). Functional enrichment of these clusters showed that they can be identified with interferon signalling pathway and translation, respectively (**Fig. 1c**). Interferon signalling genes have been previously shown to decrease translation in a number of ways^31^; here we found that with increasing immune age, this functional link between interferon activation and decrease in translation becomes even stronger (**Fig. 1d**). These findings raised the question of what driving factors might underlie the strengthened interferon–translation coupling observed with immune ageing, prompting us to search for regulators of this remodelling.

### Regulatory changes implicate miRNAs causing gut leakiness

To further explore the age-related strengthening of interferon signalling impact on suppression of translation, we next asked which genes have their expression correlated with this gene regulatory remodelling in our cohort. To do that, we employed the same sliding window principle as in our pCNA analysis. For each window, we quantified the correlation between interferon and translation clusters (**Fig. 1b** and **1d**) and the average expression of each gene. Next, we correlated these two quantities across all window positions and ranked genes according to this metric (**Fig. 2a**). Applying gene set enrichment analysis^32^ on the resulting ranked list, we identified biological processes most associated, positively or negatively, with the immune ageing-related regulatory remodelling (**Fig. 2b**). Notably, besides broad non-specific cellular processes of translation and mitochondrial respiration, we identified an enriched set of miRNA-targeted genes in the lymphocytes (**Fig. 2b**, **Supplementary Table 3**). Of note, no such result could be obtained by simply correlating gene expression with IMM-AGE across our cohort (*Methods*), further justifying our phenotype-driven correlation approach.

**Figure 2:**
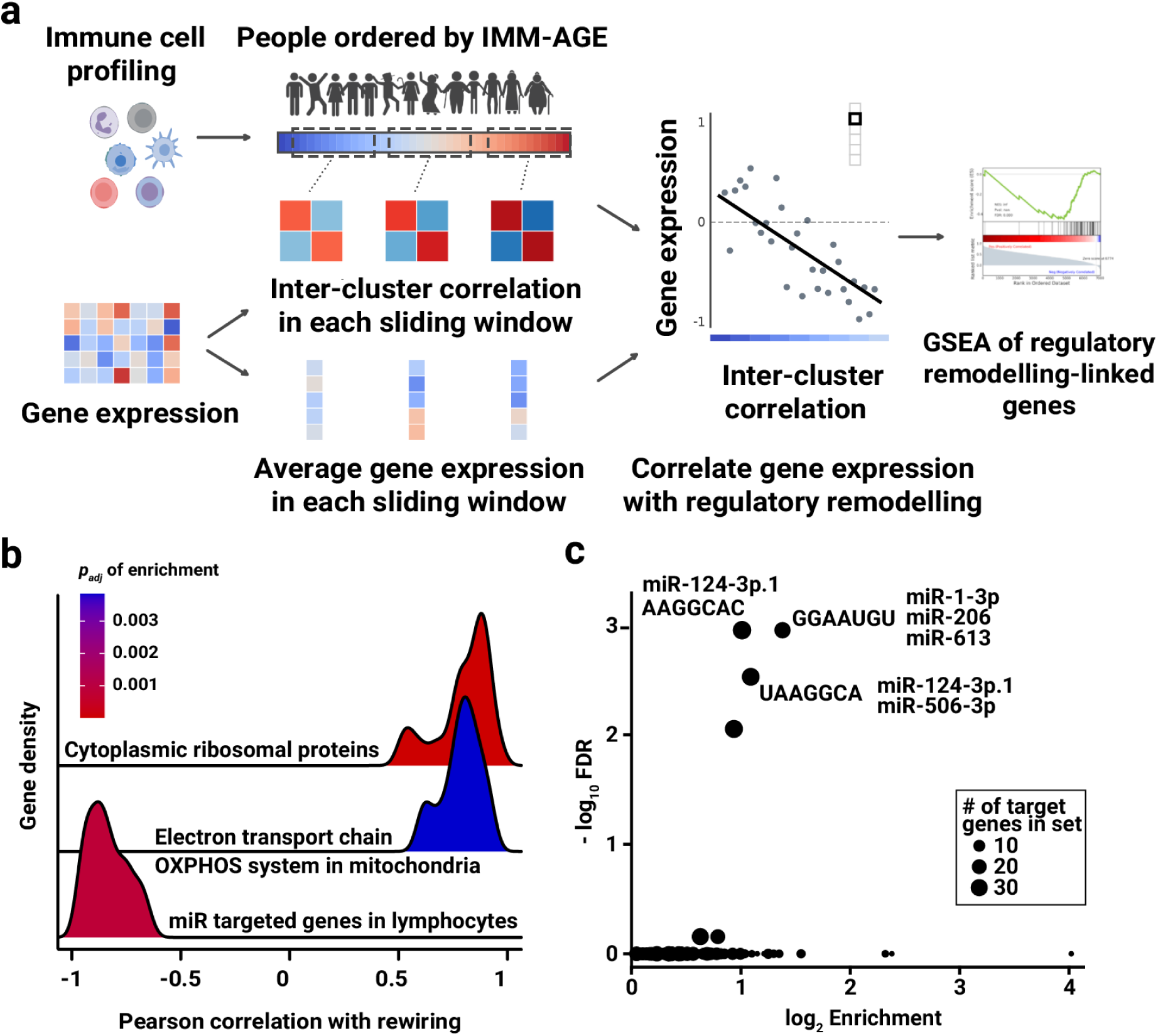
Age-dependent gene module remodelling is associated with reduced expression of miRNA regulated genes. **a**, Schematic of the strategy used for finding candidate drivers of age-associated gene regulation remodelling. Gene expression of individual genes is averaged within each position of the sliding window used in pCNA and is then correlated with translation-interferon signalling correlation (Fig. 1d right) across all window positions. The resulting list of genes ranked by these correlations is then analysed using Gene Set Enrichment Analysis (GSEA). **b**, Ridgeplot of correlations with age-dependent regulatory remodelling for genes in enriched functional groups found by GSEA. Wikipathways (WP) gene ontology was used; all terms with Benjamin-Hochberg corrected hypergeometric test *p_adj_* < 0.01 are shown. **c**, Enrichment analysis of miRNA target motifs in transcripts of genes enriched in (**b.**) belonging to *‘miR targeted genes in lymphocytes’* WP gene set using *enrichMiR*. Top 3 most significant miRNA binding motifs are labelled with the names of the corresponding miRNAs that target the motif.

To further explore the potential identity of the driver miRNA or the processes implicated, we performed a reverse search of miRNA target motifs on these genes^33^ (**Fig. 2c**). Strikingly, the two motifs with lowest FDR-values were represented by MiR-1-3p and the immune regulatory^34^ MiR-124-3p, respectively (**Fig. 2c**, **Supplementary Tables 4 and 5**), which were previously reported to synergistically damage the intestinal barrier in ageing gut, leading to increased bacterial permeability ‒ a so-called gut leakiness^35^. Since interferon signalling is known to be downstream of many pathogen sensing mechanisms^36^, including bacterial, these results together led us to hypothesise that the increased signalling through interferon pathway associated with immune ageing could be due to an increased bacterial exposure of the immune system.

### Bacterial DNA from immune cells predicts the pace of immune ageing

To further explore the hypothesis that increased bacterial exposure might drive immune ageing, we leveraged recent reports that immune cells of even healthy individuals contain measurable amounts of bacterial DNA debris^6^. We reasoned that if immune ageing is driven by increased bacterial exposure, the amount of bacterial debris in the immune cells may be associated with immune ageing. We quantified bacterial DNA extracted from PBMCs of subsets of individuals from our cohort using the same method as in the original study^6^, by performing qPCR and sequencing with primers highly specific to the bacterial ribosomal 16S RNA (**Extended Data Fig. 1 and 2**, **Supplementary Table 7**). We first tested the total bacterial DNA load in the immune cells in relation to the change in IMM-AGE score of the donors over the following year (**Fig. 3a**, *Methods*). We observed that the total bacterial DNA load was predictive of the change in immune age in a non-monotonic fashion (**Fig. 3b**). More specifically, the magnitude of excess immune ageing rate depended exponentially on the bacterial DNA load (*p* = 1.6 · 10^-5^, **Fig. 3c**). Intriguingly, for those ageing slower than average, bacterial DNA load was positively associated with immune ageing; conversely, for those ageing faster than average, bacterial DNA presence was negatively associated with immune ageing. This association was not dependent on the immune age of the subject (**Extended Data Fig. 3a-b**). No similar association was observed for the yearly IMM-AGE change before the sampling nor for the subject’s chronological or immune age at the time of sampling (**Extended Data Fig. 3c-e**).

**Figure 3:**
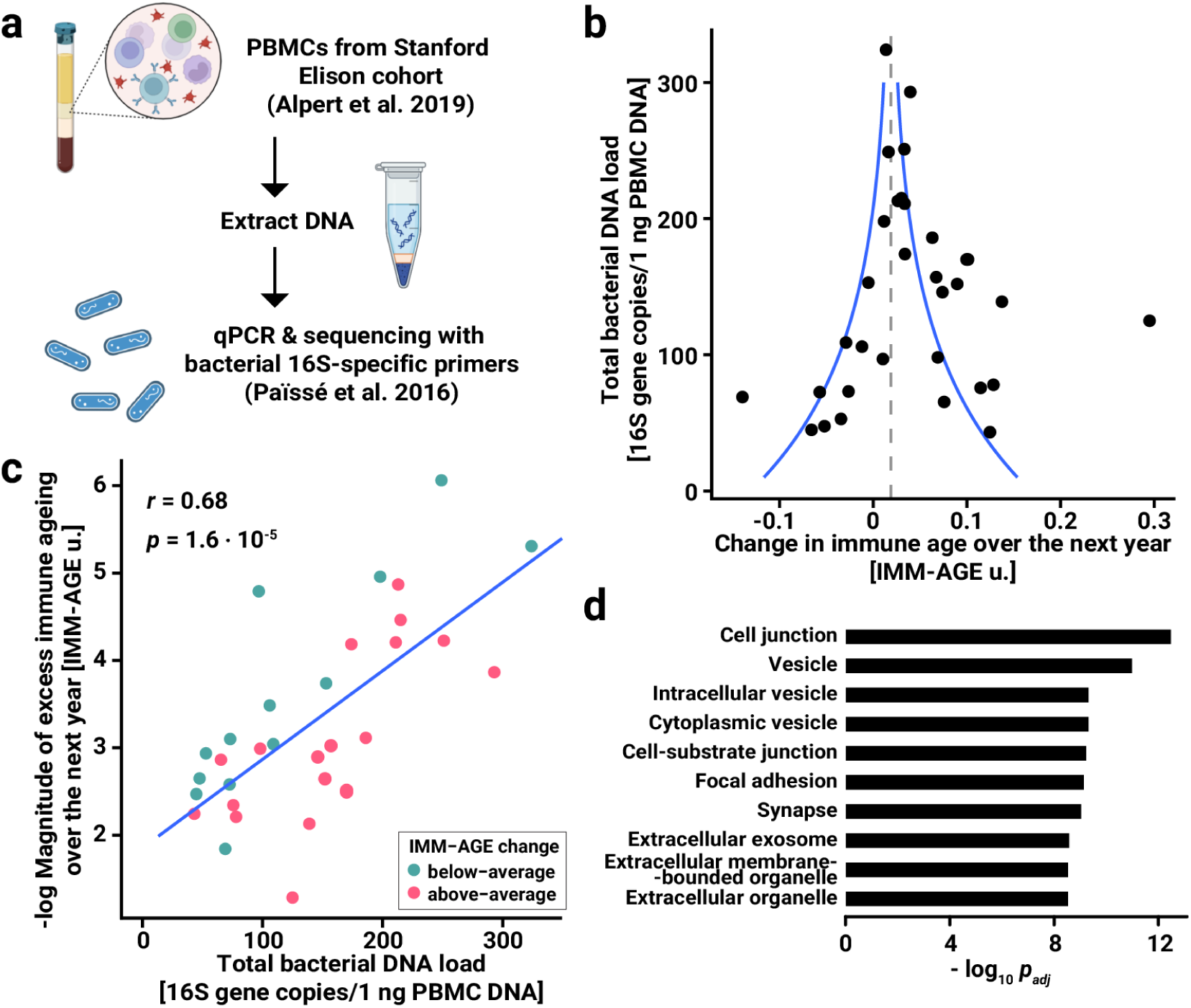
Bacterial DNA present in immune cells predicts one-year rate of immune ageing. **a**, Schematic of the experimental design. DNA from the PBMCs sampled as part of the Stanford Elison cohort was extracted and the bacterial DNA quantified using qPCR with 16S specific primers. **b**, Yearly change in IMM-AGE of a subset of Stanford-Elison cohort in relation to the total bacterial DNA load found in their PBMCs. Bacterial DNA load was determined by qPCR with bacterial 16S rRNA-specific primers on DNA extracted from PBMCs. Immune age at the time of this sampling and one year later was determined using immune cell profiles at the respective timepoints. The red dashed line represents the average yearly rate of immune ageing in the entire cohort; blue lines correspond to a fitted line in (**c.**). **c**, Exponential fit of the dependence between total bacterial DNA load from (**b.**) and the magnitude of excess immune ageing. Excess immune ageing was determined as the yearly change in IMM-AGE compared to the average yearly change in the entire cohort. *P* value and Pearson *r* are shown for the linear fit plotted in blue. **d**, Functional enrichment of differentially expressed genes between subjects with positive and negative excess yearly immune aging in Stanford Elison cohort. Samples with positive excess immune ageing over the next year were compared to those with negative excess immune ageing; 10 most significant *GO Cellular Component* terms are shown that are enriched among genes overexpressed in samples with positive excess immune ageing; no terms among under-expressed genes reached top 10 significance.

To rationalise this unusual dependence, we postulated that the presence of bacterial DNA debris in the PBMCs might be a compound readout of the opposing effects of frequency of bacterial encounters and the rate of bacterial clearance. To further investigate this hypothesis, we performed a whole blood differential gene expression analysis between all samples in our cohort with higher-than-averaging immune ageing rate as judged from a paired sample one year later, and those with similarly determined lower-than-average rate. In those ageing faster than average, we found an overexpression of genes involved in vesicle-mediated transport (**Fig. 3d**), including extracellular exosomes, potentially a part in a recently described antibacterial defense mechanism^37^. These results are broadly consistent with a scenario where a threshold level of bacterial exposure switches on a defence mechanism resulting in an increased rate of debris clearance, however, at the cost of accelerated immune ageing. More importantly, it suggests that the bacterial DNA content can provide quantitative estimates of future immune ageing over a one year period if supplemented with a gene expression readout.

We next wondered whether it could be an exposure to specific types of bacteria that is driving the immune ageing. We analysed the presence of specific bacterial taxons in the immune cells as quantified from 16S rDNA targeted sequencing (16S sequencing). We analysed the presence of specific bacterial taxons in the immune cells as quantified from 16S sequencing and their relation to immune age and its annual change. We considered both relative and absolute quantities of different bacterial phyla, orders, families and genera. After accounting for multiple hypothesis testing, only *Ralstonia* genus showed mildly significant correlation with the change in donor immune age over the next year (*p_adj_*= 0.049, **Extended Data Fig. 3f**). This suggests that it is likely the burden of the total bacterial exposure, rather than a specific bacterial taxon, that is driving the ageing of the immune system.

### Cell types increasing during immune ageing respond to bacteria

Previously, we have characterised immune ageing through quantitative changes in immune profiles, i.e. the frequencies of individual immune cell subtypes in one’s immune system^2^. Therefore, we sought to find a plausible rationale of why particular immune cell subtypes should be increasing in their frequency due to increased bacterial exposure. To address this question in an unbiased way, we subjected the immune cell subtypes that we previously identified to be associated with immune ageing to a functional analysis of their gene expression profiles. First, to generate gene expression profiles of these highly specific cell subsets, we turned to a CITE-seq atlas of PBMCs^38^ and generated pseudobulk gene expression profiles after *in silico* gating (**Fig. 4a**, **Extended Data Fig. 4 and 5**). To rank each gene for its association with increase in cell types where it is expressed, we correlated its expression in each of the cell subtypes with the relative increase of that cell subtype with age (**Fig. 4b**). Through functional gene set enrichment analysis of this ranked list, we found that the immune cell types most increasing with age are those that most express the genes involved in response to bacterial lipopolysaccharide (**Fig. 4c**; *p_adj_* = 10^-5^). This effect was not driven by a small number of cell types or genes, but rather reflects a systemic tendency of cells expressing these bacterial-response genes to increase in frequency during immune ageing (**Fig. 4d**).

**Figure 4:**
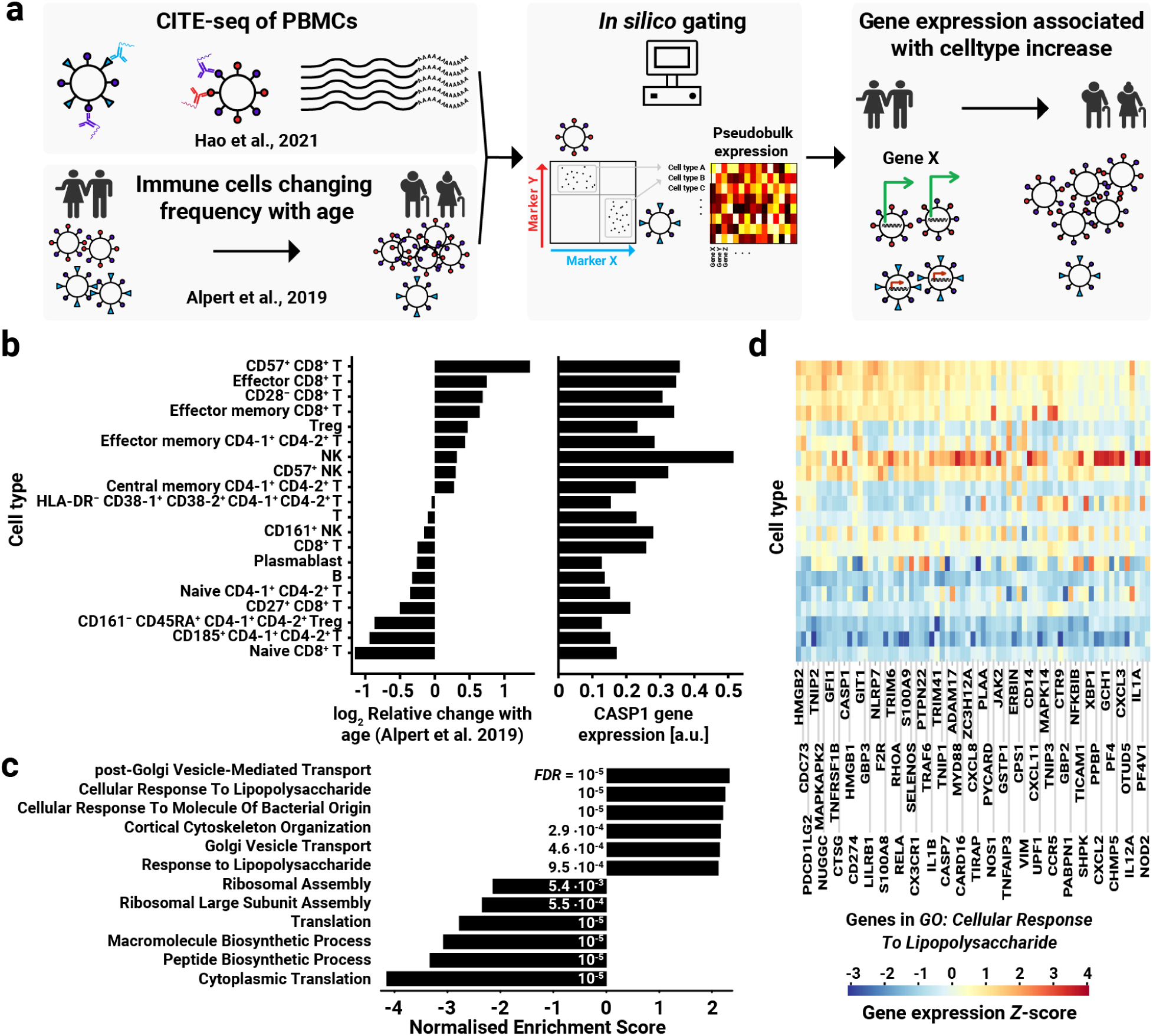
Immune cell subtypes increasing during immune ageing express bacterial response genes. **a**, Schematic of the approach. Single-cell CITE-seq data from atlas of PBMCs were gated in silico to pseudobulk gene expression in immune cell subtypes previously found associated with immune ageing (**left, middle**). Gene expression of individual genes was then associated with changes in frequencies of expressing cells across immune cell types associated with immune ageing (**right**). **b**, Example of a gene, *CASP1*, the expression of which is associated with the immune ageing-dependent change in the frequencies of immune cell subtypes. Relative change with age for each cell subset (**left**) is plotted alongside with the expression of *CASP1* in that subset (**right**). Cell subsets are ordered according to their relative change with age. **c**, GSEA of genes ranked by the Pearson correlation between their expression in respective cell types and the cell frequency change with immune ageing. GO gene ontology was used. **d**, Expression heatmap of genes from *GO:0071222* term *Cellular Response to Lipopolysaccharide*. Gene-wise *Z*-score across cell types associated with immune ageing is shown. The same cell types, and in the same order, are shown as in (**b**); genes are ordered according to the correlation of their expression in respective cell types with the age-dependent change in frequencies of these cell types; top half of the genes is shown.

## Discussion

Age-associated changes in the immune system composition in terms of specific immune cell subsets have been connected to increased mortality^2^. In this study, we devised the phenotype-driven gene correlation network analysis, a way to analyse changes in gene regulatory networks concomitant with these age-dependent changes in cell frequencies. We found remodelling of gene regulation suggestive of increasing bacterial exposure of the immune system, potentially through leaky gut. We subsequently found that the presence of bacterial DNA in immune cells as determined by 16S-sequencing in the peripheral blood mononuclear cells (PBMCs) quantitatively predicts the ageing of the immune system over the next year. This finding paves the way for a simple diagnostic marker to screen for immune systems most at risk for ageing, and given the implication of immune aging on other organs, on individuals most at risk of ageing in general. At the same time, it suggests a new research direction into antibacterial interventions in seemingly healthy people primed for decline.

It remains to be explored what processes in the human body lead to the presence of bacterial DNA in PBMCs, predictive of immune ageing. While some have suggested the presence of live bacteria in the blood of healthy people^39,40^, it is likely that most of the DNA present in PBMCs is a sentinel of what is going on elsewhere in the body, as recently demonstrated for platelets^41^, another peripheral blood component. One possibility to explain the atypical dependence of bacterial DNA content in the immune cells with respect to the direction of immune ageing is that it is a compound readout of the propensity of bacterial infections in peripheral organs and the failure of the immune system to clear them, which increases with age^42^. In this scenario, while little to no bacterial DNA present in the immune cells could indicate an overall absence of bacterial infections and thus increased prospects of good healthspan, it could also indicate an immune system that has switched to a different mode of action which fails to clear bacterial infections, heralding a rapid decline. Indeed, in model organisms, ageing has been previously proposed to follow a two-phase model, determined by gut permeability^43,44^. In the context of our results, it is likewise fitting to consider the gut microbiome as an important provenance of the immune cell exposure to bacteria during immune ageing, even more so given that immune cells primed by their exposure to the gut microbiome have been recently shown to migrate out of the gut and influence organs as distant as the brain^45,46^. Our findings thus call for additional studies where the age-dependent changes of the gut and of the immune system are investigated in conjunction^47,48^.

Our results also suggest that the changes of immune cell profiles with age are adaptive to increased bacterial threats ‒ immune cell types that increase most are those that are involved in responding to the bacterial presence. Why should then this re-calibration of the immune system towards bacterial threats be connected with immune ageing and increased mortality? One possible explanation lies in the increasingly explored trade-offs that the immune system faces ‒ evolutionary examples suggest that the immune system trades increased resistance to bacterial threats for susceptibility to other threats, such as viral ones^49^. Given that evolutionary plasticity often phenocopies the environmental plasticity^50,51^, the ageing immune system might be attuning itself to overburdening by bacterial exposure through changes in cellular proportions only to make itself more susceptible to viral or retroviral^52^ threats. Holistic studies of the immune system burdens and their consequences in terms of the immune system long-term dynamics are thus acutely needed to quantify their relative contribution to the ageing of the immune system.

## Methods

### Phenotype-driven gene correlation analysis

#### Source data

The data used in this study were obtained from the Stanford-Ellison Longitudinal Aging (SELA) cohort, a prospective observational study of generally healthy individuals, characterised longitudinally using multiple high-throughput modalities including flow cytometry, serum proteomics, and whole-blood transcriptomics^2,29,30^. The cohort has been previously used to define IMM-AGE, a metric of immune ageing based on coordinated changes in immune cell subtype frequencies^2^.

Pseudotime IMM-AGE data were calculated based on frequencies of specific immune cell subtypes as in the original study^2^; however, we only used data from years 2012-2015 that contained a harmonised antibody panel (**Supplementary Table 1**). Briefly, we normalized abundance levels before trajectory assembly. Per cell subset, we first calculated the mean and s.d. of frequency values between the 10th and 90th percentiles. We used these values to normalize cellular frequencies by subtraction of the mean and division by s.d. We applied the diffusion maps algorithm *destiny*^53^ (v.3.18.0) on the scaled cellular frequencies of the selected cell subsets that were measured in 2012-2015. The resulting diffusion pseudotime values were scaled to a range of 0-1 (**Supplementary Table 2**).

Phenotype-driven gene correlation analysis was performed using whole blood microarray gene expression data from Alpert et al. (ref. 2). First, the data were deconvolved as in the original study to account for variations in selected cell-type proportions by removing their effect on gene expression values from each gene expression profile within each year separately^54^. Next, in order to exclude batch effects that can exert profound effects in correlation analysis, only data from the years 2013-2015 that were processed in a single batch were considered (*n* = 192).

#### Correlation analysis

All samples included in the analysis were ordered according to their increasing IMM-AGE score. A sliding window of half the sample size (96) was used to calculate pairwise gene-gene Pearson correlation between all samples within the window. Shorter window sizes were also tested and gave qualitatively the same result but with more noise. For computational ease, only the 20% most-variable genes were considered (*n* = 3,858). The IMM-AGE of samples within the window was averaged using arithmetic mean. The resulting vector of IMM-AGE across all sliding window positions was correlated to gene-gene correlations across the positions of the sliding window using Pearson correlation.

#### Functional analysis of clusters of phenotype-associated regulatory remodelling

The resulting matrix of phenotype-associated gene-gene regulatory remodelling was hierarchically clustered with *single* linkage and *Canberra* distance using the *hclust* function from *WGCNA*^28^ (v.1.72-1). The clustering tree was arbitrarily cut after visual inspection, as shown in **Fig. 1b**. Functional enrichment in these clusters was determined using *ReactomePA*^55^ (v.1.42.0) gene ontology through over-representation analysis by *clusterProfiler*^56^ (v.4.6.0).

#### Candidate driver analysis

To find candidate drivers of gene regulation remodelling, for each position of the sliding window, the average pairwise correlation between genes from the two different clusters was calculated. Then, for all genes in the microarray gene expression dataset (*n* = 19,288), mean expression in each sliding was calculated and correlated to the average pairwise correlation between clusters (Pearson). Genes were ranked according to this correlation and the resulting list was subjected to gene set enrichment analysis using *Wikipathways* gene ontology through *clusterProfiler* (v.4.6.0). To test the utility of pCNA in this discovery process, we similarly performed gene set enrichment analysis on list of genes ranked by the Pearson correlation of their expression with the IMM-AGE of the donor at the time of sampling across the cohort ‒ no gene set bar ‘*Cytoplasmic ribosomal proteins*’ was significantly (BH-adjusted *p* < 0.05) enriched.

#### miRNA regulator inference

The genes driving the enrichment of the functional term *WP2004: miR targeted genes in lymphocytes* (**Supplementary Table 3**) were subjected to search of target motifs and potential upstream regulator using web-based *enrichMiR*^33^ (v.0.99.32; **Supplementary Tables 4 and 5**).

### Bacterial DNA determination in PBMCs

#### PBMCs

Peripheral blood mononuclear cells were obtained from archived samples from Stanford-Ellison longitudinal aging (SELA) cohort, a 9-year observational study previously described in detail^2,29,30^. Briefly, individuals were generally healthy at enrollment and were followed longitudinally with repeated blood draws, before and after influenza vaccination. For the present analysis, we used pre-vaccination PBMC samples from 47 individuals profiled in years 2012-2014 (year 6 to year 8 of the entire study; **Supplementary Table 6**).

#### Genomic DNA extraction

DNA was extracted from 3 million cells per sample using an optimized tissue-specific technique. Total genomic DNA was collected in a final 100μl extraction volume. Total DNA concentrations were determined by UV spectroscopy (Nanodrop®, Thermo Scientific). The mean concentration of extracted DNA was 75.5 ± 28.7 ng/ul. All the samples from the same tissue type were processed within the same DNA extraction run by the same single experimenter and were spatially and temporally isolated from the DNA extraction runs of other tissue types. Specific measures to avoid cross-contamination have been taken such as cleaning and DNA decontamination of the safety hoods in-between each extraction run.

#### qPCR quantitation

The 16S rRNA gene copies present in the samples were measured by qPCR in triplicate and normalized using a plasmid-based standard scale. The number of bacterial 16S rRNA gene copies is assessed using the *Universal 16S Real Time qPCR* workflow established by Vaiomer (Vaiomer SAS, Labège, France). Real-time PCR amplification was performed using 16S universal primers targeting the V3-V4 region of the bacterial 16S ribosomal gene (*Vaiomer universal 16S primers*). The qPCR step is performed on a VIIA 7® PCR system (Life Technologies) using Sybr Green technology and the following amplification cycles: hold stage of 10 min at 95°C, then 40 cycles of 15 sec at 95°C, 1 min at 63°C and 1 min 72°C. The absolute number of copies of 16S rRNA gene was determined by comparison with a quantitative standard curve of 16S rRNA gene plasmids generated by serial dilution of plasmid standards (*Vaiomer Universal standard plasmids*). The total number of bacterial 16S rRNA gene copies present in the samples was measured by qPCR in triplicate and normalized using a plasmid-based standard range. The specificity of all qPCR products was assessed by systematic analysis of the post-PCR dissociation curve performed between 60°C to 95°C.

#### 16S rDNA sequencing

The microbial populations present in the samples have been determined using next generation high throughput sequencing of variable regions of the 16S rRNA bacterial gene^57^. The PCR amplification was performed using 16S universal primers targeting the V3-V4 regions of the bacterial 16S ribosomal gene (Vaiomer universal 16S primers). For each sample, a sequencing library was generated by addition of sequencing adapters. The detection of the sequencing fragments was performed with the MiSeq Illumina technology using the 2 x 300 paired-end MiSeq kit V3.

#### 16S rDNA sequence analysis

The targeted metagenomic sequences from microbiota were analyzed using a FROGS bioinformatics pipeline (v.3.2.3)^58^. Briefly, after demultiplexing of the barcoded Illumina paired reads, single read sequences were cleaned and paired for each sample independently into longer fragments. The following specific filters have been applied: amplicons without the two PCR primers were removed (10% of mismatches are allowed); amplicons with at least one ambiguous nucleotides (N) were removed; the last 10 bases of reads R1 were removed; the last 50 bases of reads R2 were removed; amplicons with a length < 350 nt or a length > 500 nt were removed. Samples with less than 5000 sequences after FROGS processing were not included in the downstream analysis. Operational taxonomic units (OTUs) were produced via single-linkage clustering and taxonomic assignment was performed in order to determine community profiles. OTUs matching any of the following filtering criteria were removed: OTUs with abundance lower than 0.005% of the whole dataset abundance; OTUs identified as chimeras with *vsearch*^59^ (v.2.17); OTUs with a strong similarity (coverage and identity ≥ 80%) with the phiX (library used as a control for Illumina sequencing runs). OTUs were clustered in two passes of the *swarm*^60^ algorithm (v.3.0). The first pass was a clustering with an aggregation distance equal to 1, the second pass a clustering with an aggregation distance equal to 3.The taxonomic assignment was produced by *Blast+*^61^ (v2.10.1) with the databank *Silva*^62^ (v.138.1 Parc). Rarefaction calculations were performed using an *R* package *Vegan*^63^ (v.2.5-6), used in Python with *rpy2*. Scikit-Bio (v.0.4.2) was used for calculating α-diversity.

#### Quality controls and exclusions

To assess the potential DNA contamination from the environment and the sampling material, four freezing media samples, four replicates of the pool of supernatants and four DNA extraction negative controls were added and carried over throughout the 16S rRNA gene sequencing pipeline and assessed for difference in bacterial DNA content to the PBMC samples (**Supplementary Table 7**). There was a clear separation between PBMC samples, the DNA extraction negative controls, and the supernatant pool, as judged by the quantity of bacterial DNA.

For four PBMC samples (15-015-01-15, 15-019-01-15, 15-080-03-15 and 15-129-01-15), only reads belonging to *Pseudomonas* were detected. These samples were also the four having the highest amount of bacterial DNA in qPCR. This is consistent with either an ongoing bacterial infection present in the donor(s), or a contamination of the original samples with Pseudomonas DNA/bacteria. These samples were therefore excluded from further analysis. Seven samples (15-041-03-13, 15-048-01-14, 15-061-01-13, 15-062-01-13, 15-073-01-14, 15-125-01-15, 15-131-01-14) which amplified without issue in the qPCR, did not amplify correctly in PCR during the 16S sequencing library preparation. These samples which may contain more PCR inhibitors, were reprocessed in a new PCR step with different conditions. The seven samples were successfully amplified in the new PCR, but this introduced a contaminant (*Aeromonas* genus) corresponding to 7% to 18.8% of the reads of these 7 samples while being virtually absent from all other samples. This taxon was subtracted during the bioinformatic analysis, removing the corresponding 7% to 18.8% of the total reads in the seven samples, and between 0% and 0.006% of the total reads in the other samples. Finally, samples of PBMCs from after the vaccination were not considered in the analysis of bacterial DNA effect on immune age to exclude the effect of vaccination itself.

#### Association of bacterial DNA to immune ageing

The total bacterial load as normalised to the unit mass of DNA extracted from PBMCs was related to the IMM-AGE change over the next year by subtracting the mean yearly IMM-AGE change in the entire considered cohort (only paired samples one year apart were considered) from the IMM-AGE change data, taking an absolute value and logarithm and subsequently fitting a linear regression model to its relation to the bacterial load using *stat_regline_equation* function of *ggpubr*^64^ (v.0.6.0).

The presence of specific bacterial taxons in the immune cells as quantified from 16S sequencing (**Supplementary Tables 8-12**) was related to immune ageing in four different ways – the taxon relative frequency to current immune age, the taxon relative frequency to change in IMM-AGE over the next year, the taxon absolute abundance to current immune age and the taxon absolute abundance to change in IMM-AGE over the next year. Absolute taxon abundance was quantified by multiplying its relative abundance in the respective donor and the total 16S DNA count from the donor per unit mass DNA isolated from PBMCs. We considered bacterial taxons at the level of phylum, order, family and genus. Function *cor.test* from R package *stats* (v.4.3.1) was used to quantify Pearson correlation and the corresponding *p* value, that was subsequently Bonferroni-adjusted by multiplying with the number of taxa tested at the given level.

### Differential gene expression for 1-year immune ageing

To analyse differentially expressed genes between people with positive and negative excess immune ageing, the same gene expression dataset was used as for the pCNA (see above), but subsetted for samples that had a paired sample from the same donor with IMM-AGE pseudotime from one year later (*n* = 156). Excess immune ageing was then defined as the difference between the actual immune ageing in the next year for the given sample and the mean one-year immune ageing across this subset of samples. The sample set was then divided into two groups of positive and negative excess immune ageing and differentially expressed genes determined using *limma*^65^ (v. 3.54.1). Top 5% most overexpressed and underexpressed genes according to the moderated *t-*statistic were subjected to functional over-representation analysis with Cellular Component Gene Ontology using *ShinyGO*^66^ (v.0.85).

### Gene expression analysis of cell-types associated with immune ageing

#### Dataset Used

The dataset originates from a study on human peripheral blood mononuclear cells (PBMCs), sampled from eight donors (aged 20-49) participating in a clinical trial for HIV vaccine adjuvant and sequenced using the CITE-seq protocol^38^. The uniqueness of this dataset lies in the breadth of antibody library used for this study – the total number of antibody markers sums up to 228; in total, this dataset consists of ∼211,000 cells that were sequenced by this experimental setup. The dataset was downloaded from Gene Expression Omnibus, acc. no. GSE164378, and converted into a Python-compatible *AnnData* structure. Only donors in the control group were considered for our study (*n* = 5; ∼16,000 cells); data from the three timepoints were analysed both collectively as well as stratified by the timepoint.

#### Cell Subtype Identification

Our immune cell subtypes definitions in terms of surface markers were tailored to closely follow those used by the original IMM-AGE study^2^. We identified several surface markers that were used to identify cell subtypes associated with immune ageing in the original study, but were not present in the CITE-seq dataset. These were CD33, CD94, CXCR3, PD1 and CCR7. For our analysis, we substituted CCR7 with CD27 and excluded the cell types that were not suitably defined in our dataset, such as those that required one of the non-measured surface markers, from further analysis (**Supplementary Table 13**). To determine the expression threshold for binarisation of marker expression used assisted non-linear least squares fitting to model the bimodal distribution of expression (**Extended Data Fig. 4**). After visual inspection of the expression histograms, some markers were gated manually (**Extended Data Fig. 5**). Applying a sequential combination of logical *AND* conditions, derived from the marker composition of a specific immune cell subtype, to the antibody-derived tag (ADT) matrix, produced a boolean mask that marked the cells belonging to the queried subtype. This mask was then used for the quantification of gene expression stratified by cell subtype by pseudobulking ‒ single-cell records were aggregated by computing the mean expression value per cell subtype.

#### Relating Gene Expression to Immune Ageing Dynamics

We assessed the relationship between the pseudobulk gene expression patterns and the yearly mean frequency changes reported in the original immune ageing study^2^ using Pearson correlation. Specifically, for each gene, we correlated its expression across cell types with the logarithm of mean yearly frequency change of the cell types as reported in Alpert et al., 2019. The resulting correlation values were used to form a ranking of genes associated with age-dependent cellular dynamics, which we employed for gene set enrichment analysis (GSEA). For the implementation of the gene set enrichment analysis, we chose the module *prerank* introduced in the Python library *GSEApy*^67^. This method follows the original algorithm introduced by Subramanian et al. in their GSEA study^32^. We used *GO_Biological_Process_2023* as the gene ontology. The results of functional enrichment were largely consistent irrespective of whether data from all timepoints were merged (as shown in **Fig. 4**) or if individual timepoints in the dataset were used.

## Supporting information

All Supplementary Tables combined

## Data Availability

Raw data from 16S sequencing are available at European Nucleotide Archive, study no. PRJEB105417. Other data associated with this study are available as Supplementary Tables 1-13.

## Code Availability

The code associated with the analysis in this study is available on GitHub, https://github.com/lukacisinlab/Code-accompanying-Feldman-et-al-2026.

## Acknowledgments

We would like to thank Anne Deflisque for technical assistance, Marta Lukačišinová for fruitful discussions and critical reading of the manuscript, Timothy J. Few-Cooper for their comments on the manuscript and all members of S.S.S.O. and M.L. laboratories for stimulating discussions. This study was supported in part by NIH NIAID award PO1 No. A153559-01 (S.S.S.O.), the EU NextGenerationEU through the Recovery and Resilience Plan for Slovakia under the project No. 09I01-03-V05-00005 (M.L.) and the Slovak Research and Development Agency under the Contract no. VV-MVP-24-0271 (M.L.).

## Author Contributions

Conceptualisation: E.F., M.L., S.S.S.O.; Study design: E.F., H.T.M., B.L., N.M., M.L., S.S.S.O.; Experimentation: H.T.M., A.R.; Data analysis and investigation: E.F., M.H., A.R., B.L., M.L.; Figure design: M.H., M.L.; Writing ‒ original draft: M.L.; Writing ‒ review and editing: M.L., S.S.S.O.; Supervision: M.L., S.S.S.O.; Funding acquisition: M.L., S.S.S.O.

## Additional Information Statement

Correspondence and requests for materials should be addressed to Martin Lukačišin, and Shai S. Shen-Orr.

## Ethics Declaration

S.S.S.O. holds equity and is a consultant of CytoReason and holds an unpaid position with the Human Immunome Project; M.L. holds equity and is a CEO of Macrobian; A.R. and B.L. are employees and shareholders of Vaiomer.

## Extended Data Figures

**Extended Data Figure 1:**
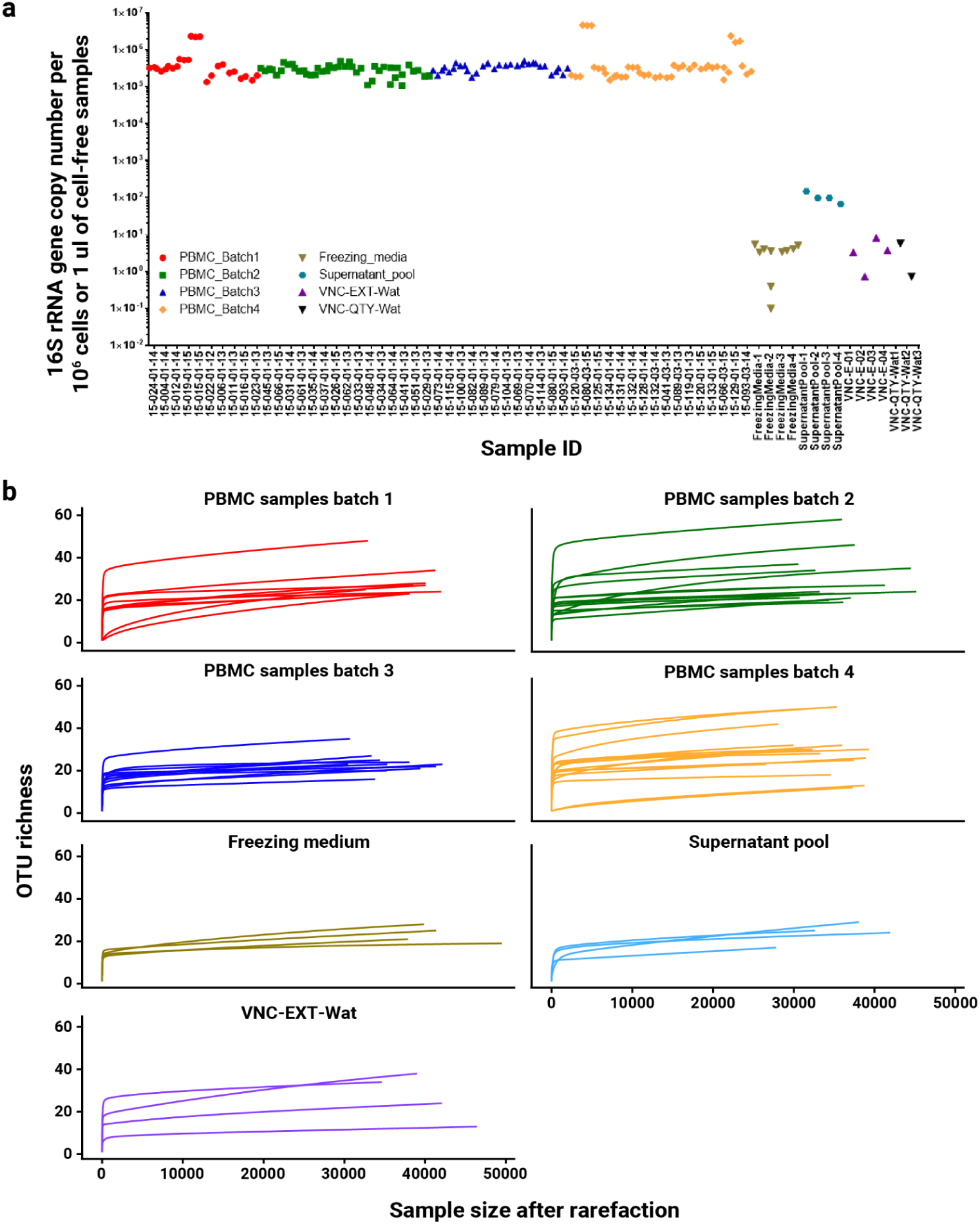
Quality controls for bacterial DNA 16S qPCR and sequencing. **a**, Bacterial DNA content detected by 16S qPCR in PBMCs samples and negative controls; for samples containing cells it is stated relative to 10^6^ PBMCs; for cell-free negative controls, it is stated by arbitrary choice relative to 1 ul of volume. Note that irrespective of this choice, the Freezing Media samples, Supernatant pool replicates and DNA extraction negative controls have a quantity of bacterial DNA measured in 16S qPCR several orders of magnitude lower than the levels found in PBMCs. **b**,Rarefaction analysis was used to assess the exhaustivity of bacterial taxa derived from the sequencing results. Rarefaction curves were created by randomly re-sampling the pool of *N* sequences multiple times and plotting the average number of OTUs found in each sample. The saturating shape of the rarefaction curves suggests that the number of reads was sufficient to capture the bacterial diversity with a good precision.

**Extended Data Figure 2:**
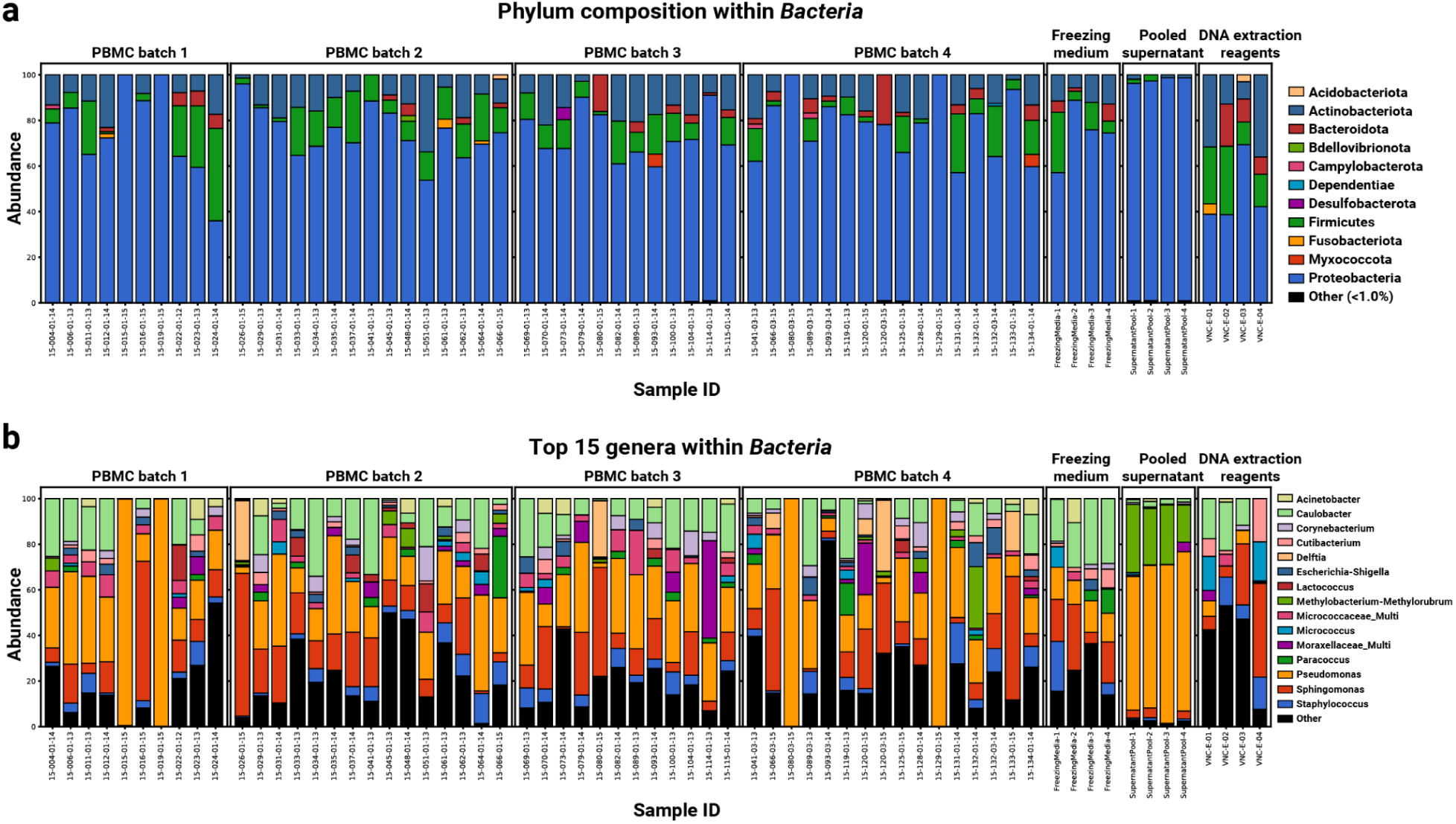
Relative abundance of taxa for different taxonomic levels. Graphical representations of the relative abundance of taxa for the taxonomic level of phylum (**a.**) and genus (**b.**) for all study samples. The most abundant taxa are identified by name in the plot, while the less abundant ones are merged into the “Other” category. The DNA extraction negative controls display variability between replicates, confirming the absence of strong contamination of the molecular biology pipeline. OTUs that are affiliated to two or more taxa are merged into a multi-affiliation (‘_Multi’) category.

**Extended Data Figure 3:**
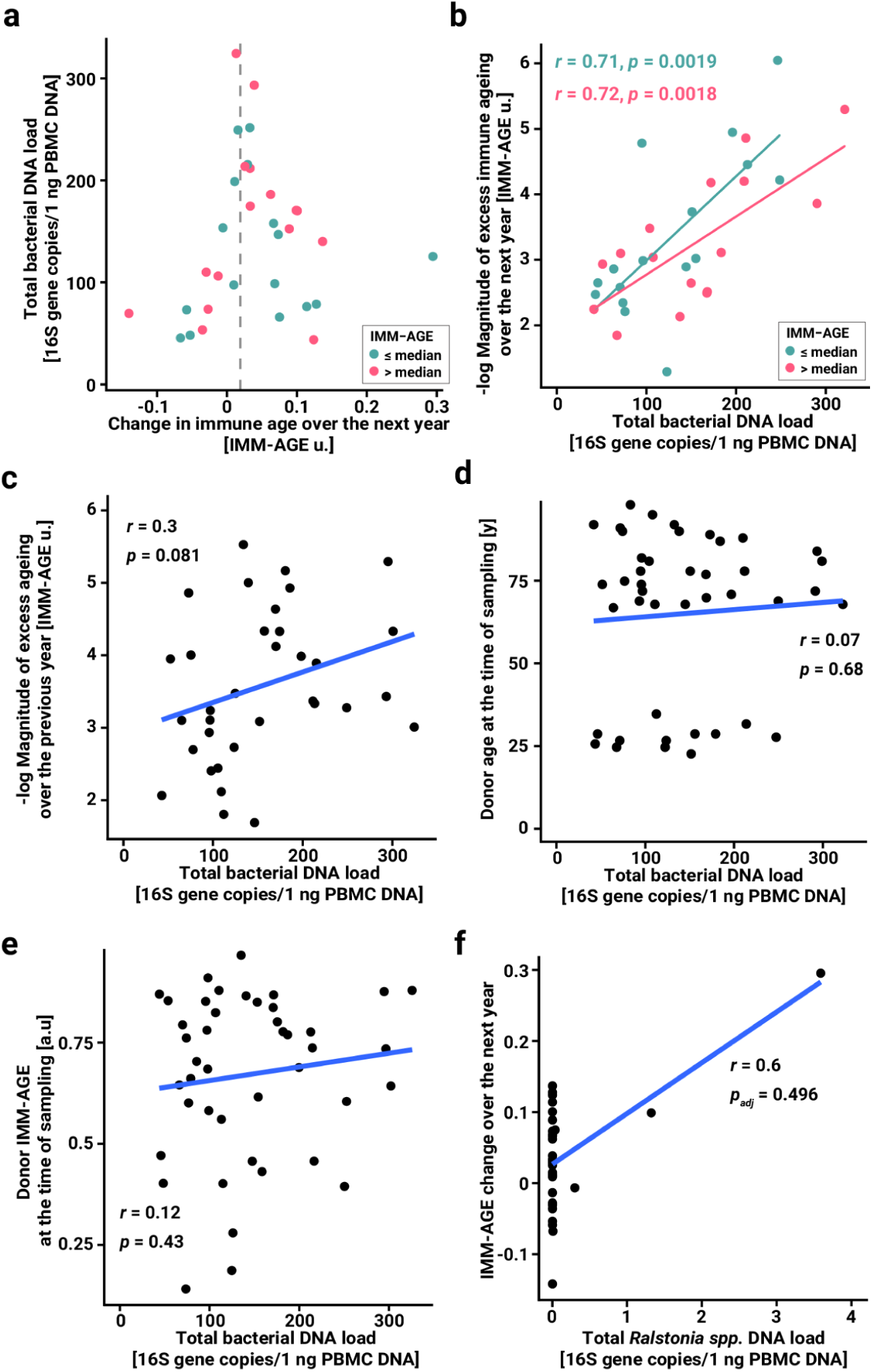
Bacterial DNA present in PBMCs is not associated with age, immune age or change in immune age over the previous year, nor is specific bacterial taxon related to immune ageing. **a-b**, As in **Fig. 3b-c**, but grouped by IMM-AGE at the timepoint of measuring bacterial DNA. **c**, Exponential fit of the dependence between total bacterial DNA load and the magnitude of excess ageing over the previous year. Excess ageing was determined as the yearly change in immune age compared to the average yearly change in the entire cohort. *P* value and Pearson *r* are shown for the linear fit plotted in blue. **d**, Linear fit for the dependence of total bacterial DNA load and donor chronological age at the time of sampling. **e**, Linear fit for the dependence of total bacterial DNA load and donor immune age at the time of sampling. **f**, Linear fit for the dependence between the absolute abundance of 16S DNA from *Ralstonia* genus and the donor’s immune age change over the next year. No other taxon showed correlation with immune age or 1-year change in immune age with *p_adj_* < 0.05. Bonferroni correction was applied to the analysis at each taxonomic level separately (phylum, order, family, genus).

**Extended Data Figure 4:**
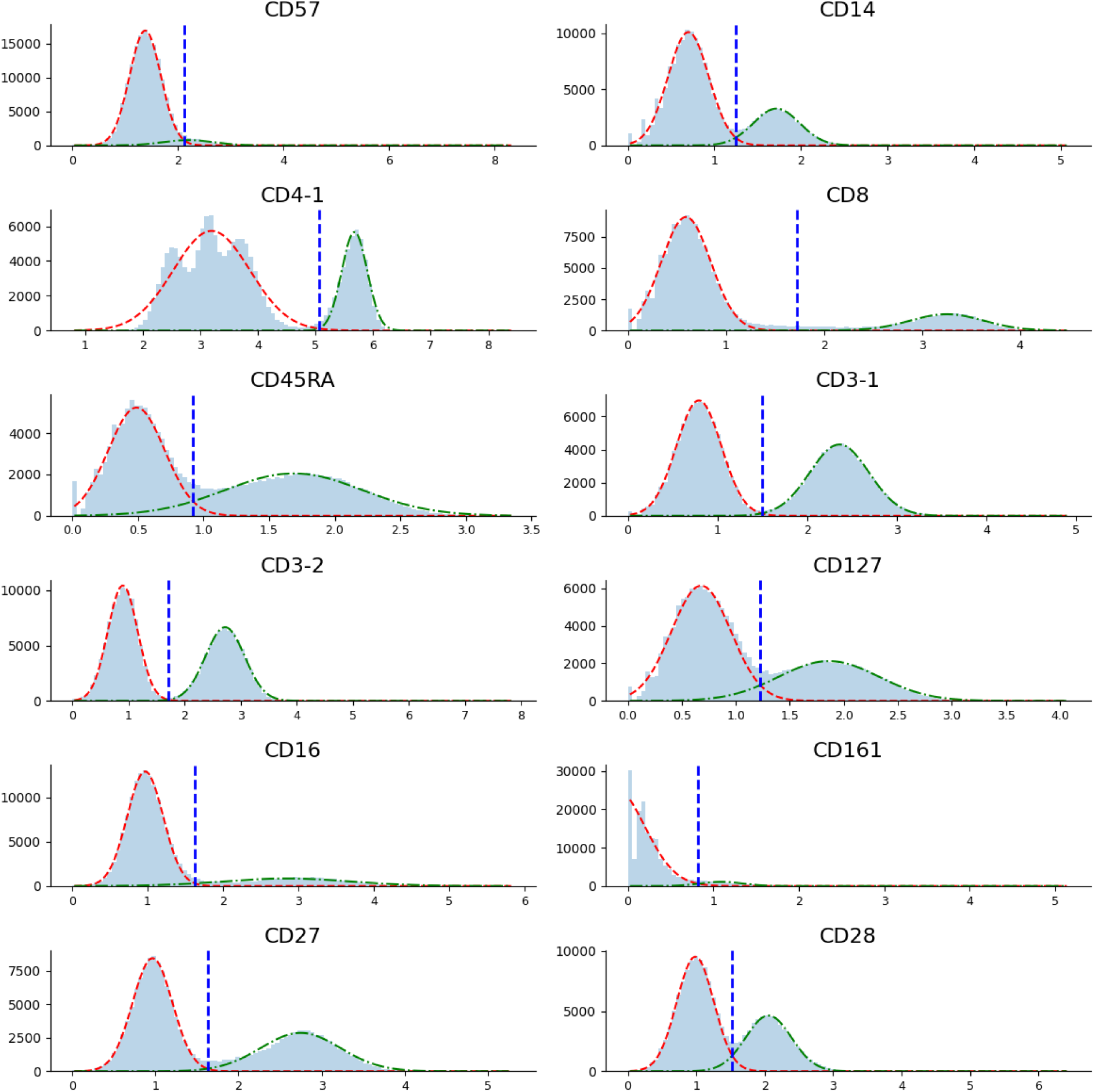
Automated thresholding for immune cell subtyping. Visualisation of thresholds produced by automatic approximation (see *Methods*). Each of the 12 histograms represent the distribution of the respective marker expression (a.u.) across all cells. The dashed red line represents the first fitted Gaussian, the green line the second one. The blue dashed line represents the estimated threshold defined by the intersection of said Gaussians.

**Extended Data Figure 5:**
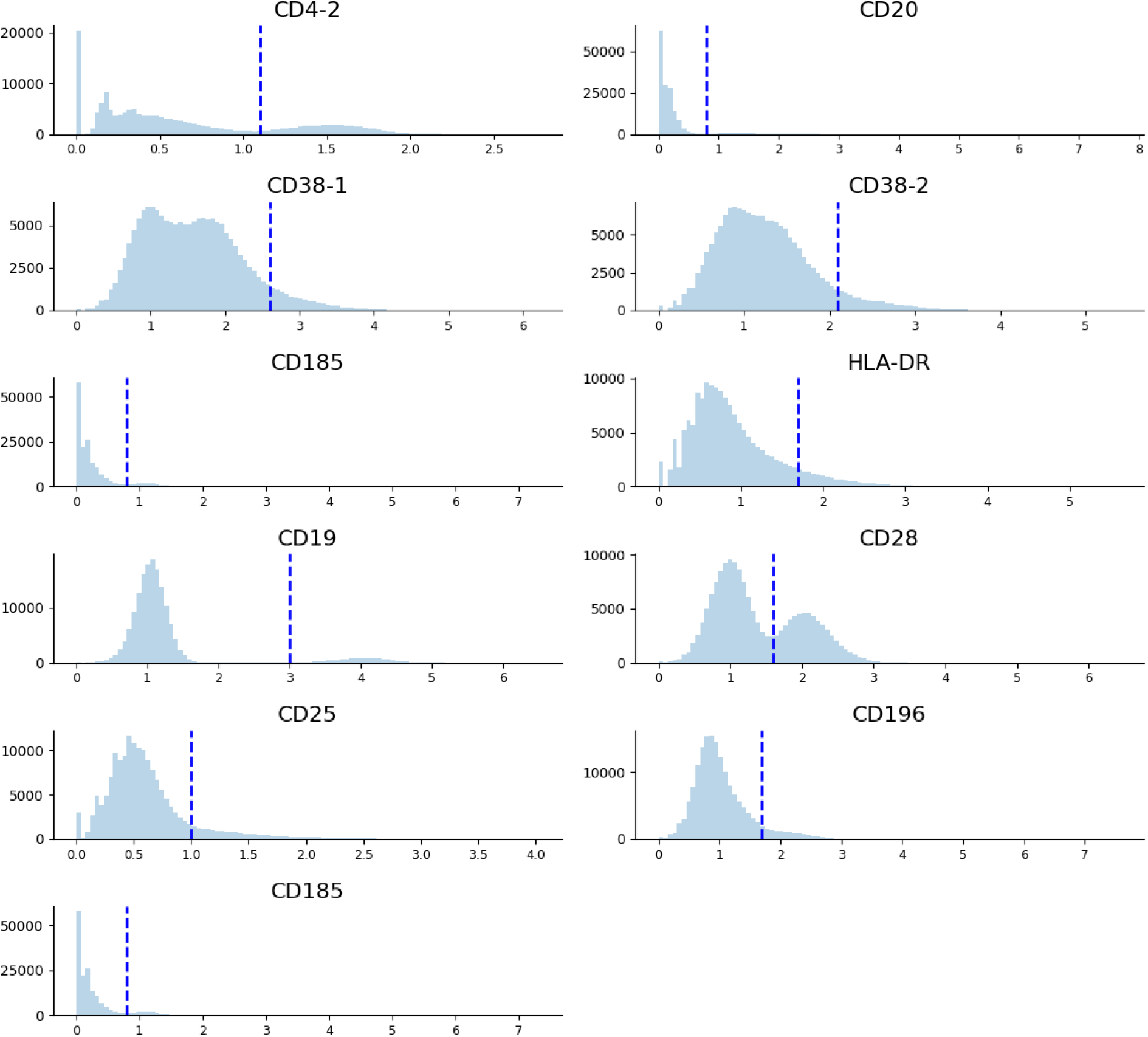
Manual thresholding for immune cell subtyping. Visualization of manually defined thresholds. Each of the 11 histograms represent the distribution of the respective marker expression (a.u.) across all cells. The blue dashed line represents the threshold defined by an educated choice.

## Supplementary Tables Captions

**Supplementary Table 1: Adjusted immune cell profiles from Alpert et al. 2019 considered in this study.**

**Supplementary Table 2: Metadata and IMM-AGE pseudotime for samples from Alpert et al. 2019 considered in this study.**

**Supplementary Table 3: Genes driving the enrichment of ‘miR targeted genes in lymphocytes’ WP gene set**

**Supplementary Table 4: Background genes considered in the gene *enrichMiR* enrichment analysis**

**Supplementary Table 5: Motifs from *enrichMiR* analysis enriched with *FDR* < 0.05**

**Supplementary Table 6: Metadata of PBMC samples used for bacterial 16S sequencing.**

**Supplementary Table 7: Bacterial 16S content in PBMC samples determined by qPCR.**

**Supplementary Table 8: 16S sequencing read counts for individual operational taxonomic units (OTUs).**

**Supplementary Table 9: 16S sequencing read fractions corresponding to individual bacterial phyla.**

**Supplementary Table 10: 16S sequencing read fractions corresponding to individual bacterial orders.**

**Supplementary Table 11: 16S sequencing read fractions corresponding to individual bacterial families.**

**Supplementary Table 12: 16S sequencing read fractions corresponding to individual bacterial genera.**

**Supplementary Table 13: Surface markers used for gating of CITE-seq data from Hao et al.**

